# High quality genome assembly and annotation (v1) of the eukaryotic terrestrial microalga *Coccomyxa viridis* SAG 216-4

**DOI:** 10.1101/2023.07.11.548521

**Authors:** Anton Kraege, Edgar A. Chavarro-Carrero, Nadège Guiglielmoni, Eva Schnell, Joseph Kirangwa, Stefanie Heilmann-Heimbach, Kerstin Becker, Karl Köhrer, WGGC Team, DeRGA Community, Philipp Schiffer, Bart P. H. J. Thomma, Hanna Rovenich

## Abstract

Unicellular green algae of the genus *Coccomyxa* are recognized for their worldwide distribution and ecological versatility. Most species described to date live in close association with various host species, such as in lichen associations. However, little is known about the molecular mechanisms that drive such symbiotic lifestyles. We generated a high-quality genome assembly for the lichen photobiont *Coccomyxa viridis* SAG 216-4 (formerly *C. mucigena*). Using long-read PacBio HiFi and Oxford Nanopore Technologies in combination with chromatin conformation capture (Hi-C) sequencing, we assembled the genome into 21 scaffolds with a total length of 50.9 Mb, an N50 of 2.7 Mb and a BUSCO score of 98.6%. While 19 scaffolds represent full-length nuclear chromosomes, two additional scaffolds represent the mitochondrial and plastid genomes. Transcriptome-guided gene annotation resulted in the identification of 13,557 protein-coding genes, of which 68% have annotated PFAM domains and 962 are predicted to be secreted.

## Introduction

Green algae are photosynthesizing eukaryotic organisms that differ greatly in terms of morphology and colonize a large variety of aquatic and terrestrial habitats. Phylogenetically, green algae form a paraphyletic group that has recently been proposed to comprise three lineages including the Prasinodermophyta in addition to the Chlorophyta and Streptophyta (Li et al., 2020). This new phylum diverged before the split of the Chlorophyta and Streptophyta that occurred between 1,000 and 700 million years ago (Morris et al., 2018). While the streptophyte lineage encompasses charophyte green algae as well as land plants, the chlorophyte lineage consists of 7 prasinophyte classes, which gave rise to 4 phycoplast-containing core chlorophyte classes (Chlorodendrophyceae, Trebouxiophyceae, Ulvophyceae, Chlorophyceae) with one independent sister class (Pedinophyceae) (Leliaert et al., 2012; Marin, 2012).

The *Coccomyxa* genus is represented by coccoid unicellular green algae that belong to the class of Trebouxiophyceae. Morphologically, *Coccomyxa* spp. are characterized by irregular elliptical to globular cells that range from 6–14 x 3–6 μm in size, with a single parietal chloroplast lacking pyrenoids and the absence of flagellate stages (Schmidle, 1901). Members of this genus are found in freshwater, marine, and various terrestrial habitats where they occur free-living or in symbioses with diverse hosts (Darienko et al., 2015; Gustavs et al., 2017; Malavasi et al., 2016). Several *Coccomyxa* species establish stable, mutualistic associations with fungi that result in the formation of complex three-dimensional architectures, known as lichens (Faluaburu et al., 2019; Gustavs et al., 2017; Jaag, 1933; Yahr et al., 2015; Zoller and Lutzoni, 2003). Others associate with vascular plants or lichens as endo- or epiphytes, respectively (Cao et al., 2018a; Cao et al., 2018b; Tagirdzhanova et al., 2023; Trémouillaux-Guiller et al., 2002), and frequently occur on the bark of trees (Kulichovà et al., 2014; Štifterovà and Neustupa, 2015) where they may interact with other microbes. One novel species was recently found in association with carnivorous plants, even though the nature of this relationship remains unclear (Sciuto et al., 2019). Besides, *Coccomyxa* also establishes parasitic interactions with different mollusk species affecting their filtration ability and reproduction (Gray et al., 1999; Sokolnikova et al., 2016; Sokolnikova et al., 2022; Vaschenko et al., 2013).

Despite this ecological versatility, little is known about the molecular mechanisms that determine the various symbiotic lifestyles in *Coccomyxa*. One short read-based genome is available for *C. subellipsoidea* C-169 that was isolated on Antarctica where it occurred on dried algal peat (Blanc et al., 2012), whereas another high-quality genome has recently been made available for a non-symbiotic strain of *C. viridis* that was isolated from a lichen thallus (Tagirdzhanova et al., 2023). For *Coccomyxa* sp. Obi, LA000219 and SUA001 chromosome-, scaffold- and contig-level assemblies are available on sNCBI, respectively, as well as two metagenome-assembled genomes of *C. subellipsoidea*. To facilitate the study of *Coccomyxa* symbiont-associated traits and their evolutionary origin, we here present the generation of a high-quality chromosome-scale assembly of the phycobiont *C. mucigena* SAG 216-4 using long-read PacBio HiFi and Oxford Nanopore Technology (ONT) combined with Hi-C and RNA sequencing. Recent SSU and ITS rDNA sequencing-based re-evaluations of the *Coccomyxa* phylogeny placed the SAG 216-4 isolate in the clade of *C. viridis* (Darienko et al., 2015; Malavasi et al., 2016). Hence, this isolate will be referred to as *C. viridis* here and data have been deposited under the corresponding Taxonomy ID.

## Materials & Methods

### Sample information

*Coccomyxa viridis* (formerly *Coccomyxa mucigena*) SAG 216-4 was ordered from the Culture Collection of Algae at the Georg-August-University Göttingen (*Sammlung von Algenkulturen der Universität Göttingen*, international acronym SAG), Germany. The stock culture was reactivated in liquid modified Waris-H growth medium (McFadden and Melkonian, 1986) with soil extract and 3x vitamins (0.15 nM vitamin B12, 4.1 nM biotin, 0.3 μM thiamine-HCl, 0.8 nM niacinamide), and maintained through regular medium replacement. Cultures were grown at ∼ 15 μmol photons m^-2^ s^-1^ (fluorescent light tubes: L36W/640i energy saver cool white and L58W/956 BioLux, Osram, Munich, Germany) in a 14/10 h light/dark cycle at 20°C.

### DNA *and* RNA extraction

Cells of a 7-week-old *C. viridis* culture were harvested over 0.8 μm cellulose nitrate filters (Sartorius, Göttingen, Germany) using a vacuum pump. Material was collected with a spatula, snap-frozen and ground in liquid nitrogen using mortar and pestle. The ground material was used for genomic DNA extraction with the RSC Plant DNA Kit (Promega, Madison, WI, USA) using the Maxwell® RSC device according to manufacturer’s instructions. To prevent shearing of long DNA fragments, centrifugation was carried out at 10,000 *g* during sample preparation. Following DNA extraction, DNA fragments <10,000 bp were removed using the SRE XS kit (Circulomics, Baltimore, MD, USA) according to manufacturer’s instructions. DNA quantity and quality were assessed using the Nanodrop 2000 spectrometer and Qubit 4 fluorometer with the dsDNA BR assay kit (Invitrogen, Carlsbad, CA, USA), and integrity was confirmed by gel electrophoresis. High-molecular weight DNA was stored at 4°C.

For total RNA extraction, algal cells were collected from a dense nine-day-old culture and ground in liquid nitrogen using mortar and pestle. RNA was extracted with the Maxwell RSC Plant RNA kit (Promega, Madison, WI, USA) using the Maxwell® RSC device according to manufacturer’s instructions. RNA quality and quantity was determined using the Nanodrop 2000 and stored at -80°C.

### Pacific Biosciences High-Fidelity (PacBio HiFi) sequencing

HiFi libraries were prepared with the Express 2.0 Template kit (Pacific Biosciences, Menlo Park, CA, USA) and sequenced on a Sequel II/Sequel IIe instrument with 30h movie time. HiFi reads were generated using SMRT Link (v10; (Pacific Biosciences, Menlo Park, CA, USA) with default parameters.

### Oxford Nanopore Technologies (ONT) sequencing

Library preparation with the Rapid Sequencing Kit (SQK-626 RAD004) was performed with ∼400 ng HMW DNA according to manufacturer’s instructions (Oxford Nanopore Technologies, Oxford, UK). The sample was loaded onto an R9.4.1 flow cell in a minION Mk1B device (Oxford Nanopore Technologies, Oxford, UK), which was run for 24 h. Subsequent base calling was performed using Guppy (version 630 3.1.3; Oxford Nanopore Technologies, Oxford, UK). Adapter sequences were removed using Porechop (version 0.2.4 with default settings) (Wick, 2018), and the reads were self-corrected and trimmed using Canu (version 1.8) (Koren et al., 2017).

### Chromosome conformation capture (Hi-C) and sequencing

*C. viridis* cells were cross-linked in 3% formaldehyde for 1 hour at room temperature. The reaction was quenched with glycine at a final concentration of 250 mM. Cells were collected by centrifugation at 16,000 *g* for 10 min. Pellets were flash-frozen in liquid nitrogen and ground using mortar and pestle. Hi-C libraries were prepared using the Arima-HiC+ kit (Arima Genomics, Carlsbad, CA, USA) according to manufacturer’s instructions, and subsequently paired-end (2x150 bp) sequenced on a NovaSeq 6000 instrument (Illumina, San Diego, CA, USA).

### RNA sequencing

Library preparation for full-length mRNASeq was performed using the NEB Ultra II Directional RNA Library Prep with NEBNext Poly(A) mRNA Magenetic Isolation Module and 500 ng total RNA as starting material, except for W-RNA Lplaty, where library prep was based on 100 ng total RNA as starting material. Sequencing was performed on an Illumina NovaSeq 6000 device with 2x150 bp paired-end sequencing protocol and >50 M reads per sample.

### Genome assembly

PacBio HiFi reads were assembled using Raven (v1.8.1) (Vaser and Šikić, 2021) with default settings. Hi-C reads were mapped onto this assembly with Juicer (v2.0) using the “assembly” option to skip the post-processing steps and generate the merged_nodups.txt file (Durand et al., 2016b). For the juicer pipeline, restriction site maps were generated using the *Dpn*II (GATC) and *Hin*fI (GANTC) restriction site profile and the assembly was indexed with BWA index (v0.7.17-r1188) (Li and Durbin, 2009), and used to polish the assembly using 3d-dna (v180922) (Dudchenko et al., 2017). Afterwards, Juicebox (v1.11.08) was used to manually curate the genome assembly by splitting contigs and rearranging them according to the Hi-C pattern (Durand et al., 2016a). Contigs were merged to scaffolds according to the Hi-C map and Ns were introduced between contigs within scaffolds, gaps between contigs were removed and contigs were merged. Subsequently, ONT reads were mapped to the assembly using Minimap2 (v2.24-r1122) and Samtools (v1.10) and mapped reads were visualized in Integrative Genome Viewer (v2.11.2) (Danecek et al., 2021; Li, 2021; Robinson et al., 2011). Whenever gaps between contigs were spanned by at least five reads with a mapping quality of 30, the contigs were fused in the assembly.

Potential telomeres were identified using tapestry (v1.0.0) with “AACCCT” as telomere sequence (Davey et al., 2020). To check for potential contaminations, Blobtools (v1.1.1) and BLAST (v2.13.0+) were used to create a Blobplot including taxonomic annotation at genus level (Camacho et al., 2009; Laetsch and Blaxter, 2017). To check completeness of the assembly and retrieve ploidy information, kat comp from the Kmer Analysis Toolkit (v2.4.2) was used, and results were visualized using the kat plot spectra-cn function with the -x 800 option to extend the x-axis (Mapleson et al., 2016). Genome synteny to the closest sequenced relative *C. subellipsoidea* C-169 was determined using Mummer3 (Blanc et al., 2012; Kurtz et al., 2004). In detail, the two assemblies were first aligned using Nucmer, followed by a filtering step with Delta-filter using the many-to-many option (-m). Finally, the alignment was visualized with Mummerplot.

### Annotation

To annotate repetitive elements in the nuclear genome, a database of simple repeats was created with RepeatModeler (v2.0.3) that was expanded with transposable elements (TE) from the TransposonUltimate resonaTE (v1.0) pipeline (Flynn et al., 2020; Riehl et al., 2022). This pipeline uses multiple tools for TE prediction and combines the prediction output. For the prediction of TEs in *Coccomyxa viridis* helitronScanner, ltrHarvest, mitefind, mitetracker, RepeatModeler, RepeatMasker, sinefind, tirvish, transposonPSI and NCBICDD1000 were used within TransposonUltimate resonaTE and TEs that were predicted by at least two tools were added to the database. TEclass (v2.1.3) was used for classification (Abrusán et al., 2009). To softmask the genome and obtain statistics on the total TE and repetitive element content in the genome, RepeatMasker (v4.1.2-p1)(Smit et al., 2012) was used with excln option to exclude Ns in the masking.

Gene annotation in the nuclear genome was performed making use of RNA sequencing data. To this end, the genome was indexed, and reads were mapped with HiSat2 (v2.2.1) using default settings (Kim et al., 2019). Afterwards, BRAKER1 (v2.1.6) was used for transcriptome-guided gene prediction based on the RNA sequencing data with default settings (Hoff et al., 2016). To generate protein and coding sequence files the Braker output was transformed with Gffread (v0.12.7) (Pertea and Pertea, 2020). PFAM domain annotation was performed with InterProScan (v5.61) (Paysan-Lafosse et al., 2023). To estimate the number of secreted proteins, SignalP (v6.0) was run in the slow-sequential mode on the annotated proteins (Teufel et al., 2022). Finally, BUSCO (v5.3.2) was run with the Chlorophyta database (chlorophyta_odb10) to estimate the completeness of the gene annotation (Manni et al., 2021). The circos plot visualization of the annotation was created with R (v4.2.0) and Circilize (v0.4.14) (Gu et al., 2014). All software and tools used for the genome assembly and annotation are summarized in Table S1.

Organelle genomes were annotated separately. Scaffolds were identified as organelle genomes based on their lower GC content and smaller size. The mitochondrial genome was annotated using MFannot (Lang et al., 2023) as well as GeSeq (Tillich et al., 2017) and the annotation was combined within the GeSeq platform. The plastid genome was annotated using GeSeq alone. The annotations were visualized using the OGDraw webserver (Greiner et al., 2019).

## Results

The version 1 genome of *C. viridis* was assembled from 32.2 Gbp of PacBio HiFi reads with a mean read length of 15 kb, 0.95 Gbp Nanopore reads with a mean read length of 8.8 kb and 15 million pairs of Hi-C seq data. The PacBio HiFi reads were first assembled using Raven (Vaser and Šikić, 2021), yielding 27 contigs. These contigs were scaffolded and manually curated using Hi-C data (Dudchenko et al., 2017; Durand et al., 2016a; Durand et al., 2016b; Li and Durbin, 2009). To close the remaining gaps between contigs within scaffolds, ONT reads were mapped onto the assembly (Danecek et al., 2021; Li, 2021) and gaps that were spanned by at least 5 ONT reads with a mapping quality >30 were manually closed, finally resulting in 21 scaffolds consisting of 26 contigs with a total length of 50.9 Mb and an N50 of 2.7 Mb (Figure 1, Table 1). Using Tapestry (Davey et al., 2020), telomeric regions ([AACCCT]*n*) were identified at both ends of nine of the 21 scaffolds (≥5 repeats) (Figure 1a), suggesting that these represent full-length chromosomes, which was confirmed by Hi-C analysis (Figure 1b). Additionally, the Hi-C contact map indicated centromeres for some of the chromosomes. However, the determination of exact centromere locations on all chromosomes will require ChIP-seq analysis and CenH3 mapping. While Tapestry detected telomeric sequences at only one end of eight other scaffolds and none for scaffold 18 and 19, the Hi-C map points towards the presence of telomeric repeats at both ends of all scaffolds 1-19 (Figure 1b), suggesting that the v1 assembly contains 19 full-length chromosomes that compose the nuclear genome. Scaffolds 20 and 21 were considerably shorter with ∼162 kb and ∼70 kb and displayed a markedly lower GC content at 41-42% (Figure 1a), suggesting that these scaffolds represent the chloroplast and mitochondrial genomes, respectively. BLAST analyses confirmed the presence of plastid and mitochondrial genes on the respective scaffolds, and the overall scaffold lengths corresponded with the sizes of the plastid and mitochondrial genomes of *Coccomyxa subellipsoidea* C-169 with 175 kb and 65 kb, respectively (Blanc et al., 2012). Full annotation of scaffolds 20 and 21 showed that they indeed represent chloroplast and mitochondrial genomes, respectively (Figure 2).

**Figure 1.**
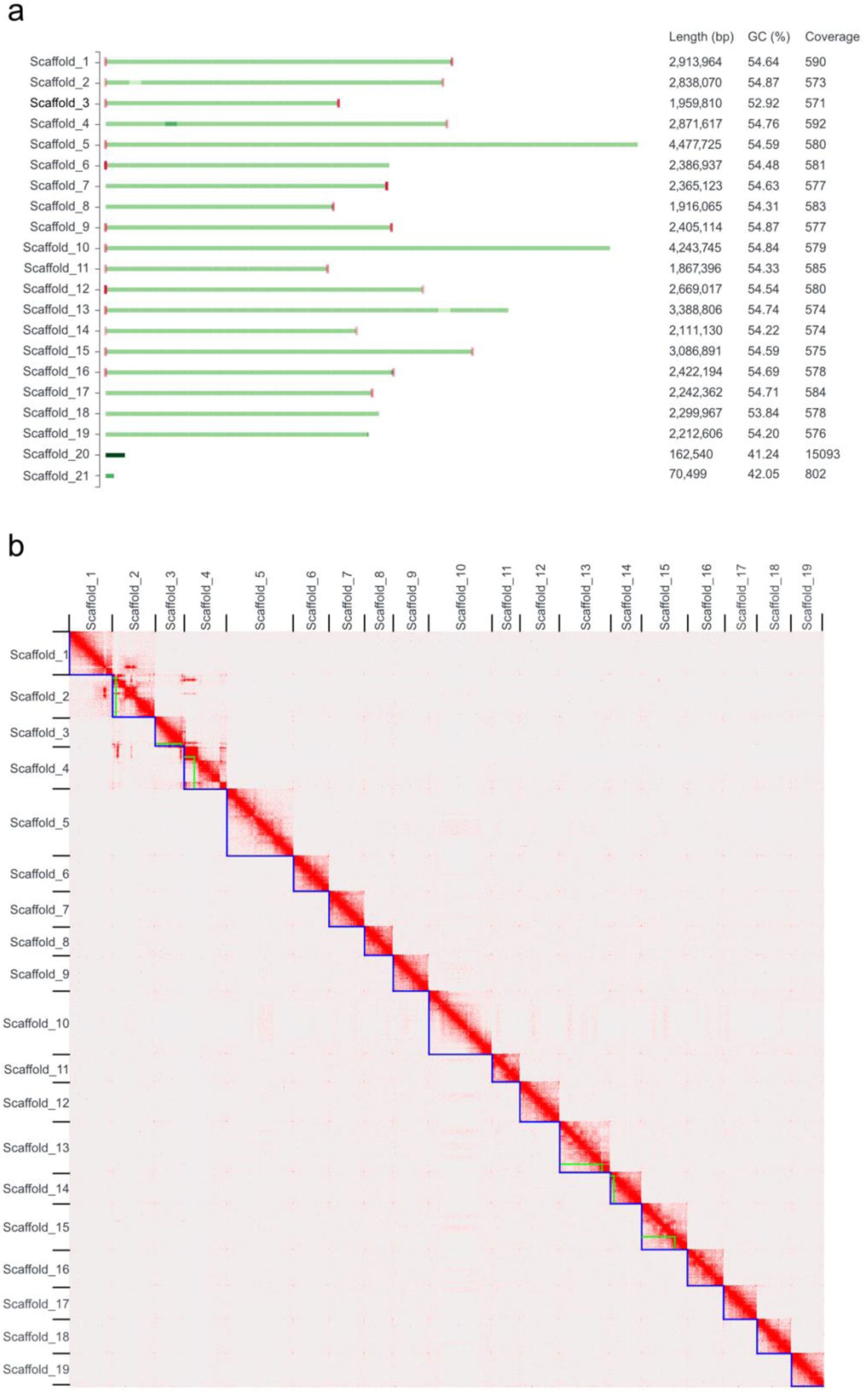
Genome assembly of *Coccomyxa viridis* SAG 216-4. (a) An overview of the *C. viridis* genome assembly depicts chromosome-scale scaffolds. Green bars indicate scaffold sizes and red bars represent telomeres. Variations in color intensities correlate with read coverage. Read coverage per scaffold is determined by mapping PacBio HiFi reads onto the assembly. Scaffolds 20 and 21 were identified as chloroplast and mitochondrial genomes based on size and low GC contents, and BLAST analyses. (b) Hi-C contact map showing interaction frequencies between regions in the nuclear genome of *Coccomyxa viridis*. Scaffolds are framed by blue lines while contigs within scaffolds are depicted in green.

**Figure 2.**
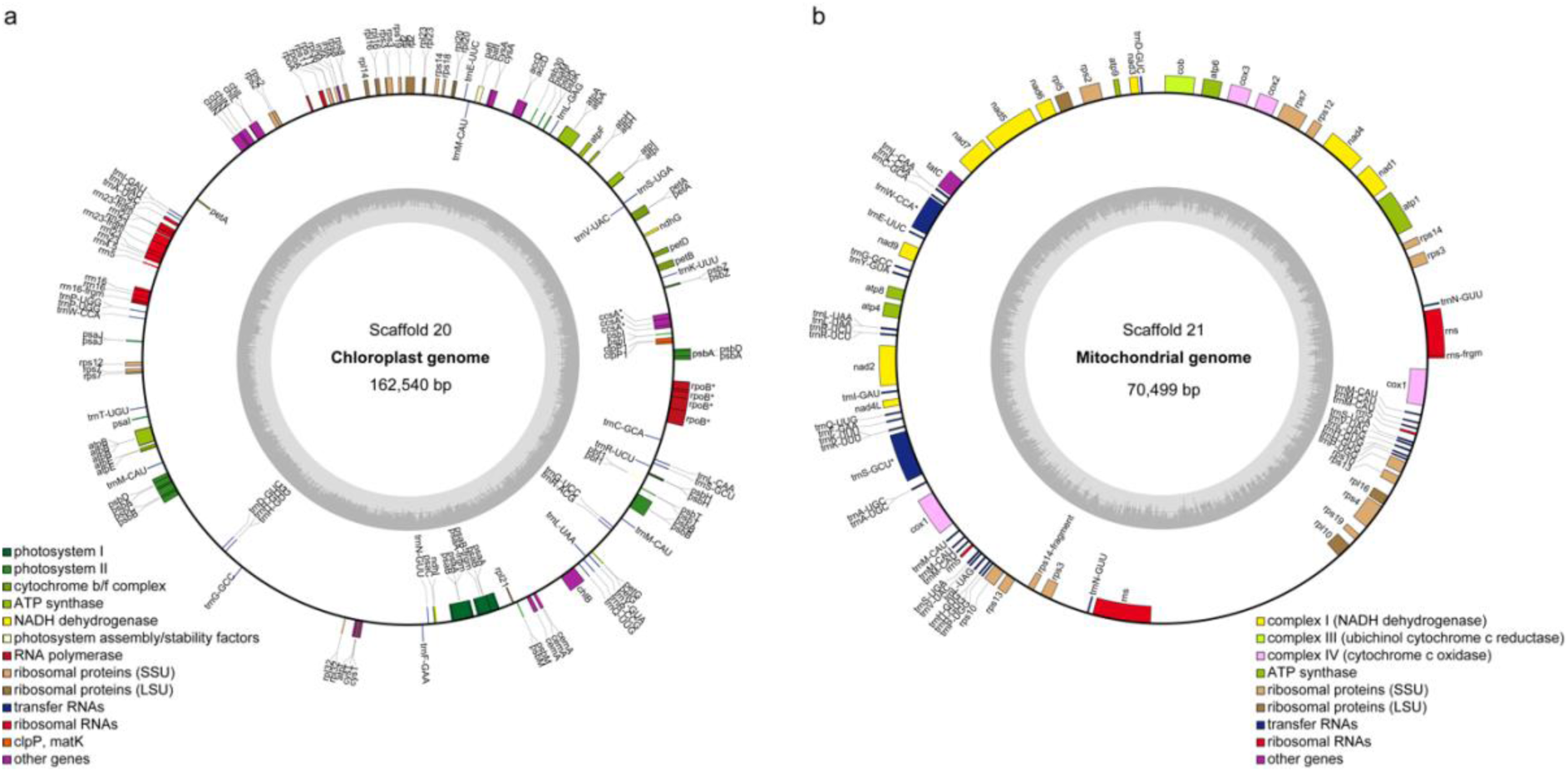
Scaffolds 20 and 21 represent the plastid and mitochondrial genomes of *C. viridis* SAG 216-4. Gene maps of the chloroplast (a) and mitochondrial (b) genomes. The inner circles indicate the GC content and mapped genes are shown on the outer circles. Genes that are transcribed clockwise are placed inside the outer circles, and genes that are transcribed counterclockwise at the outside of the outer circles.

**Table 1.** Genome features of *C. viridis* SAG 216-4 including the mitochondrial and plastid genomes.

| Assembly ID | *C. viridis* SAG 216-4 genomes |
| --- | --- |
| Total length (bp) | 50,911,578 |
| No. of contigs | 27 |
| No. of scaffolds | 21 |
| Longest scaffold (bp) | 4,477,725 |
| N50 (bp) | 2,669,017 |
| L50 | 8 |
| GC content (%) | 54.5 |

To rule out the presence of contaminants, the assembly and PacBio HiFi raw reads were used to produce a Blobplot (Camacho et al., 2009; Laetsch and Blaxter, 2017), which indicates that 98.76% of the reads match only the *Coccomyxa* genus (Figure 3) and, consequently, that the original sample was free of contaminating organisms. Finally, a KAT analysis showed a single peak of k-mer multiplicity based on HiFi reads that were represented once in the assembly (Figure 4) (Mapleson et al., 2016), indicative of a high-quality, haploid genome.

**Figure 3.**
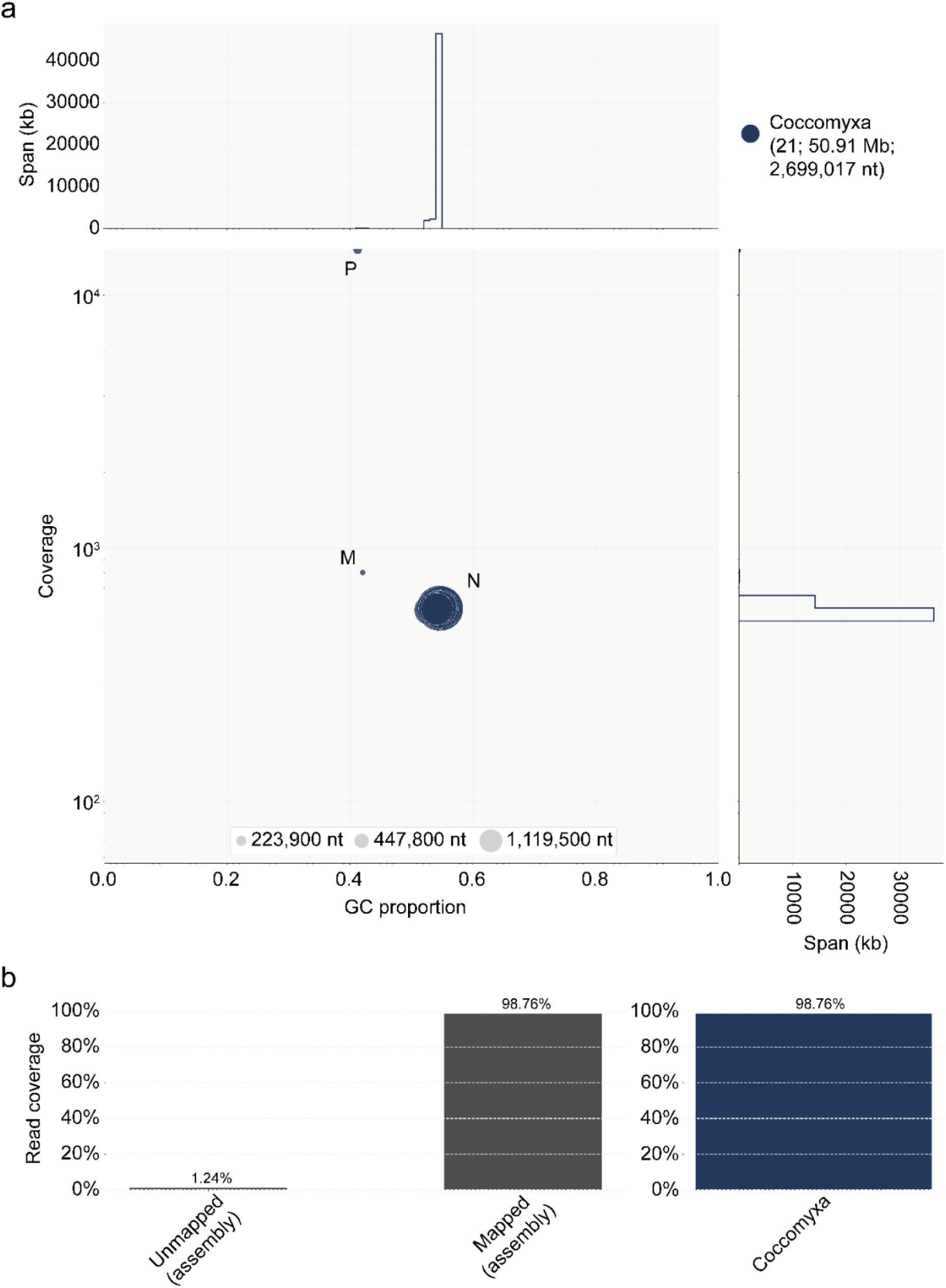
Taxonomic annotation indicates absence of contaminations in the genome assembly. (b) Taxon-annotated GC coverage scatter plot (Blobplot) of the contigs from the genome assembly shows that all scaffolds are taxon-annotated as *Coccomyxa* and all scaffolds that belong to the nuclear genome have similar GC contents (∼54%). The GC content of the mitochondrial and plastid genomes are considerably lower (∼41%). (b) In total 98.76% of the reads can be mapped onto the assembly and are therefore classified as *Coccomyxa* reads.

**Figure 4.**
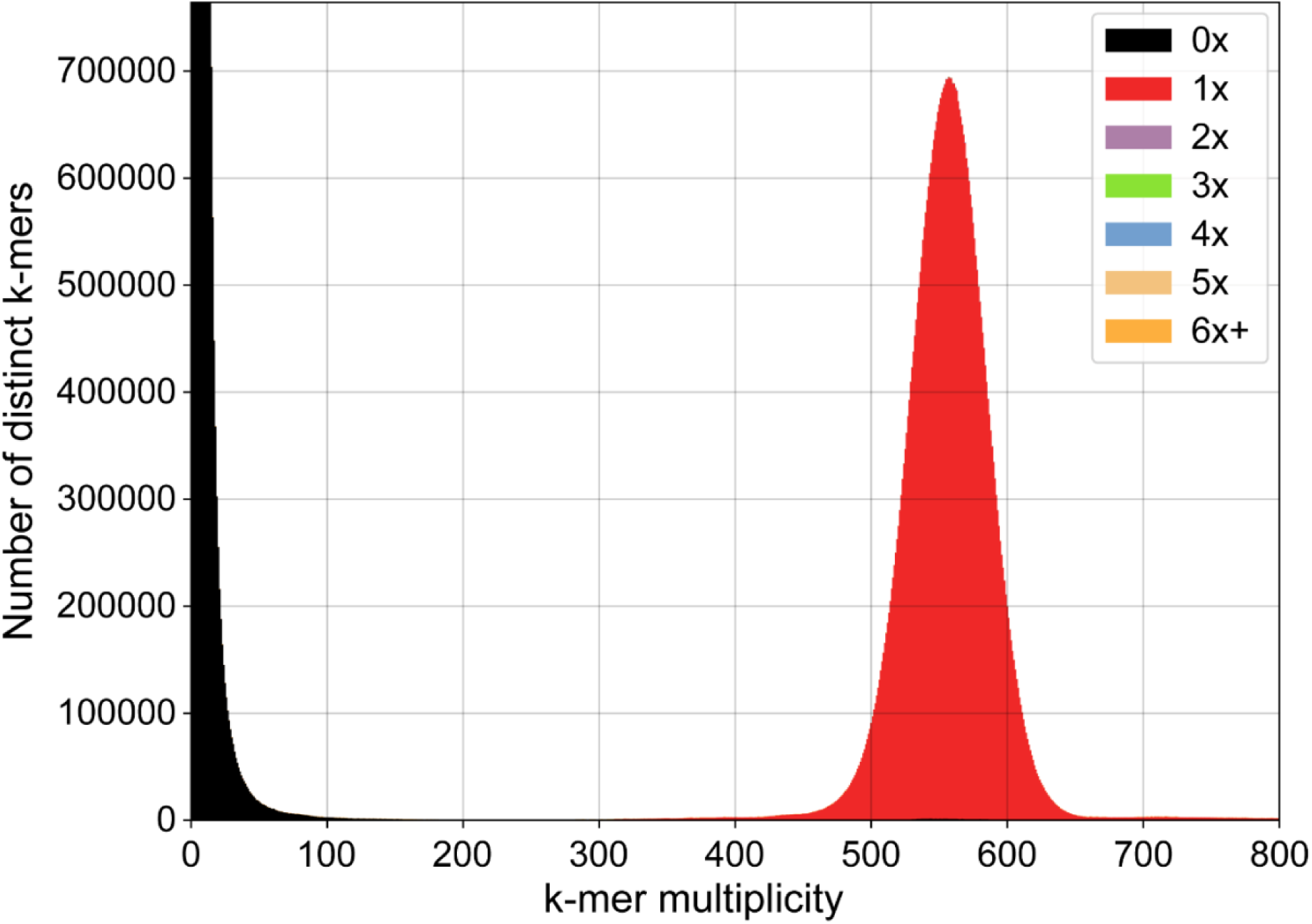
The *Coccomyxa viridis* SAG 216-4 genome is haploid. The KAT specra-cn plot depicts the 27-mer multiplicity of the PacBio HiFi reads against the genome assembly. Black areas under the peaks represent k-mers present in the reads but absent from the assembly, colored peaks indicate k-mers that are present once to multiple times in the assembly. The single red peak in the KAT specra-cn plot suggests that *Coccomyxa viridis* has a haploid genome, while the black peak at low multiplicity shows that the assembly is highly complete and that all reads are represented in the assembly.

To annotate the nuclear genome, we first assessed the presence of repetitive elements. In total, we found 8.9% of the genome to be repetitive (Table 2), comparable to the 7.2% of repetitive sequences found in the genome of *C. supellipsoidea* C-169 (Blanc et al., 2012). These 8.9% repetitive elements were annotated as either simple repeats (2.3%) or transposable elements (6.6%). Of the transposable elements, 36% were annotated as retrotransposons and 64% as DNA transposons. The distribution of the repetitive elements was even across the genome with only a few repeat-rich regions (Figure 5). Next, we aimed to produce a high-quality genome annotation using RNA sequencing data. In total 13,557 genes were annotated with an average length of 3.1 kb (Table 2). The amount of alternative splicing in the genome is predicted to be very low, given the average of one transcript per gene model. To confirm the actual amount of alternative splicing, however, further analyses will be required. Of the 13,557 genes, 68% have annotated PFAM domains and 962 are predicted to carry a signal peptide for secretion. A total of 1,489 (98.6 %) complete gene models among 1,519 conserved Benchmarking Universal Single-Copy Orthologs (BUSCO) (Manni et al., 2021) in the chlorophyta_odb10 database were identified (Table 2), suggesting a highly complete genome annotation.

**Figure 5.**
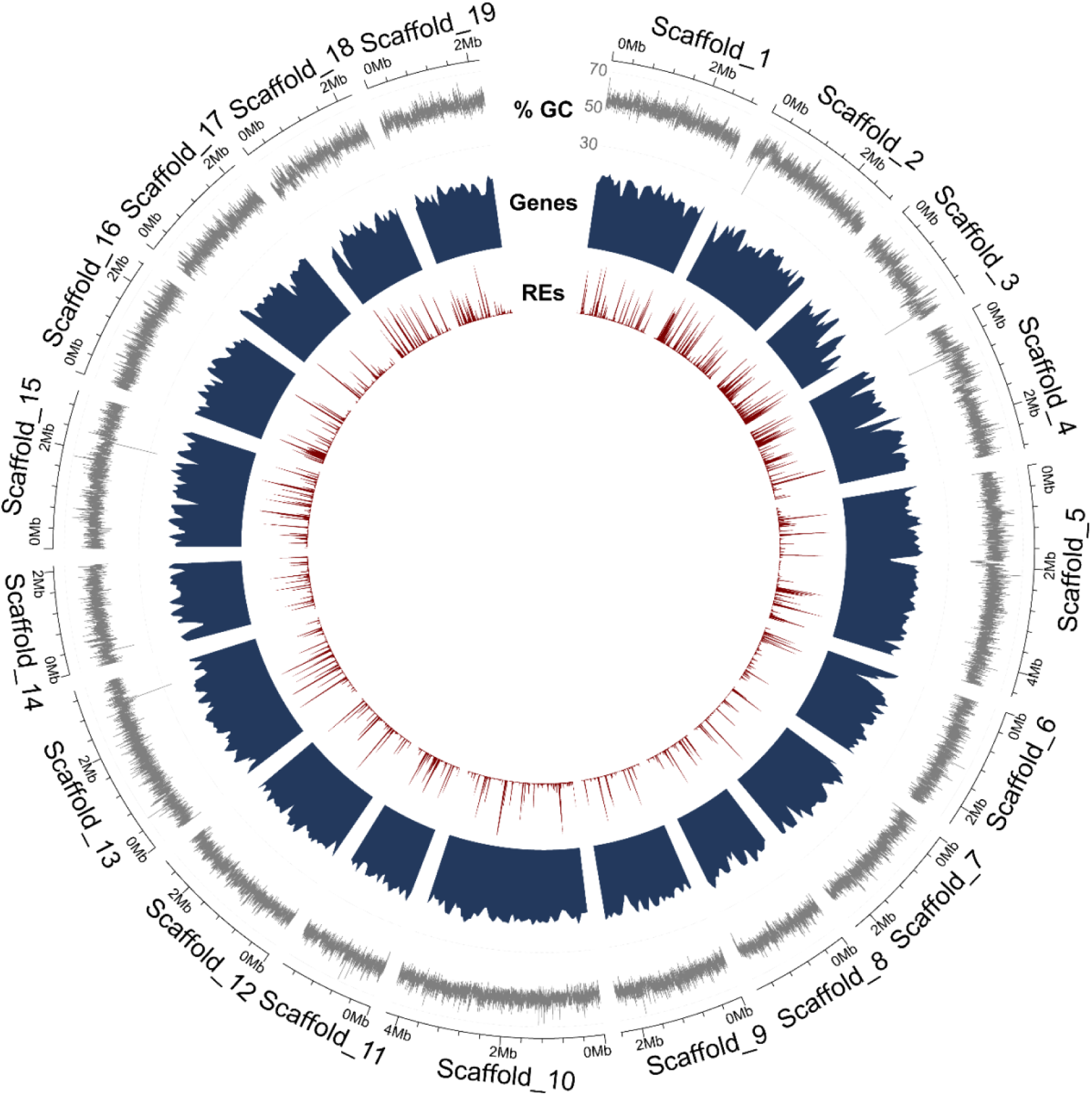
Circos plot summarizing the nuclear genome annotation of *Coccomyxa viridis* SAG 216-4. From outside to inside the tracks display: GC content (over 1-kb windows), gene density (blue) and repetitive element density (red).

**Table 2.** Annotation features of the *C. viridis* SAG 216-4 nuclear genome.

| **Genome annotation** |  |
| --- | --- |
| Repeat content (%) | 8.85 |
| Retrotransposons | 2.4 |
| DNA transposons | 4.2 |
| Simple repeats | 2.25 |
| No. gene models | 13,557 |
| Average gene length (bp) | 3146 |
| No. exons | 122,978 |
| Average no. exons per gene model | 9 |
| Average exon length (bp) | 158 |
| No. transcripts | 14,024 |
| Average no. transcripts/gene model | 1 |
| No. gene models <200 bp length | 0 |
| No. proteins with $\geq 1$ PFAM domain | 9205 |
| No. proteins with signal peptide | 962 |
| **BUSCO (chlorophyta_odb10)** | **C: 98.6% [S: 82.5%, D: 16.1%], F: 0.1%, M: 1.3%, N: 1519** |

Until recently, the taxonomic classification and definition of *Coccomyxa* species was based on environmentally variable morphological and cytological characteristics. This classification was reviewed based on the phylogenetic analyses of nuclear SSU and ITS rDNA sequences, which resulted in the definition of 27 currently recognized *Coccomyxa* species (Darienko et al., 2015; Malavasi et al., 2016). Dot plot analysis of the high-quality genome assembly of *C. viridis* SAG216-4 with the assembly of the most closely related sequenced relative *C. subellipsoidea* C-169 revealed a lack of synteny since the few identified orthologous sequences were < 1 kb and, therefore, do not represent full-length genes (Figure 6a, Table 2). This lack of synteny was no technical artifact since the *C. viridis* assembly could be fully aligned to itself (Figure 6b), and BLAST analyses with five out of six non-identical ITS sequences identified in the *C. viridis* SAG 216-4 assembly confirmed its species identity. A comparison of the assembly of *C. subellipsoidea* C-169 to that of *Chlorella variabilis* (Chlorophyte, *Trebouxiophyceae*) has previously identified few syntenic regions which displayed poor gene collinearity (Blanc et al., 2012). Future studies will help to clarify whether the absence of synteny between *C. viridis* and *C. subellipsoidea* is due to the quality of the available assemblies or whether it has biological implications.

**Figure 6.**
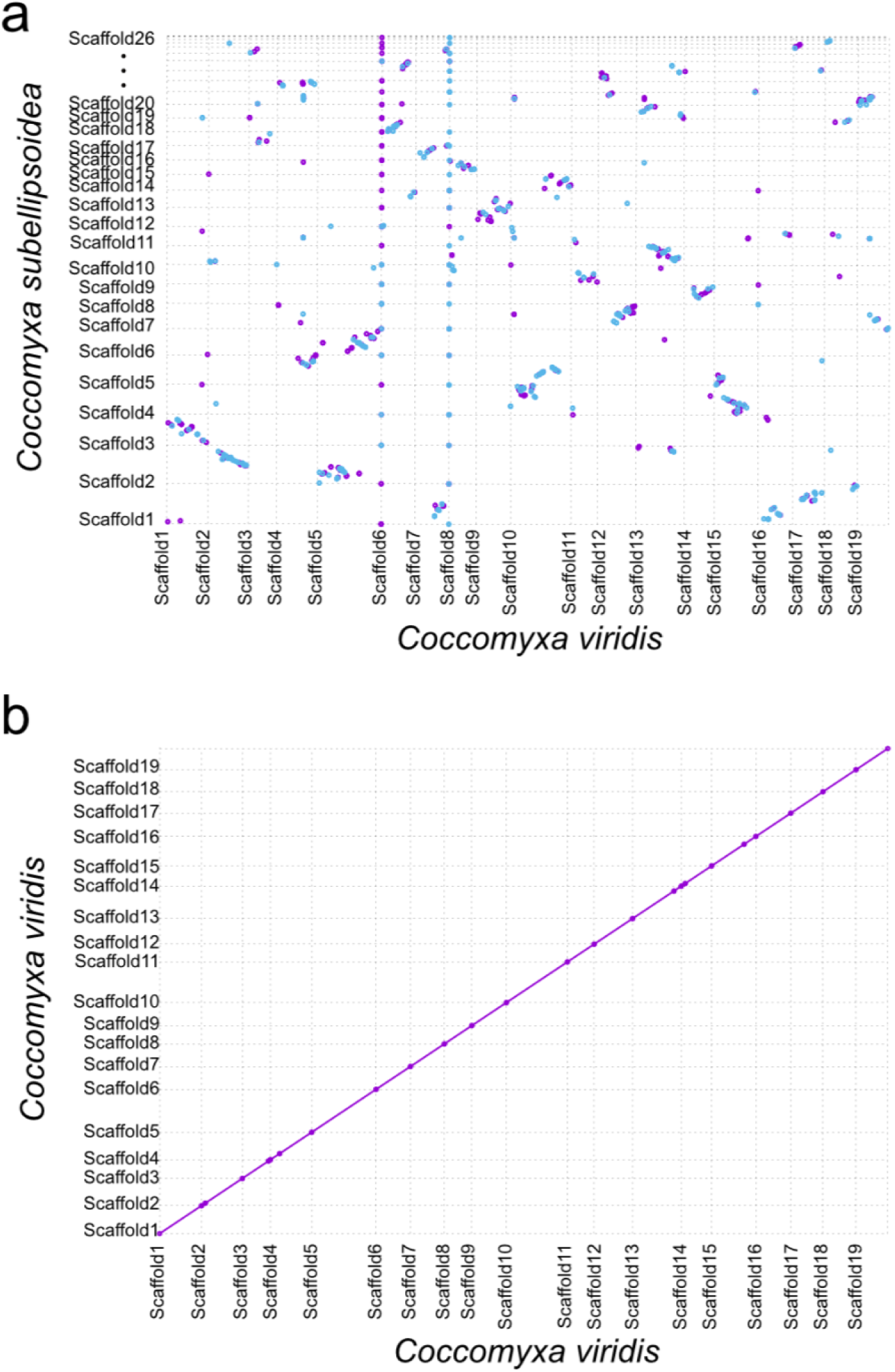
No synteny detected between related *Coccomyxa* species. (a) Dot plot of orthologous sequences in the genome assemblies of *C. viridis* SAG 216-4 and *C. subellipsoidea* C-169. Violet and blue dots represent orthologous sequences on same and opposite strands, respectively. Dot sizes does not correlate with the length of the sequences they represent, which were all < 1 kb. The width of each box corresponds to the length (bp) of the respective scaffold. (b) Dot plot of the genome assembly of *C. viridis* SAG216-4 against itself.

## Data availability

Data for *C. viridis* SAG 216-4 with the ToLID ucCocViri1 is available via the European Nucleotide Archive (ENA) under the study accession number PRJNA1054215. Fastqc reports of raw data can be found in (Kraege et al., 2023).

## Acknowledgements

This genome project is part of the DeRGA pilot study (https://www.erga-biodiversity.eu/pilot-project) and we acknowledge coordination and support by Dr. Astrid Böhne and Prof. Dr. Ann-Marie Waldvogel. Development of the pilot ERGA Data Portal (https://portal.erga-biodiversity.eu/) was funded by the European Molecular Biology Laboratory. We thank the West German Genome Center (WGGC) and the Cologne Center for Genomics (CCG) for library generation and quality control as well as their support for PacBio Hifi, Hi-C and RNA sequencing.

---

**Table S1.** Summary of bioinformatics tools used for genome assembly and annotation.

| Assembly |  | Annotation |  |
| --- | --- | --- | --- |
| Tool | Version | Tool | Version |
| Raven | v1.8.1 | RepeatModeler | v2.0.3 |
| Juicer | v2.0 | TransposonUltimate | v1.0 |
| BWA | v0.7.17-r1188 | TEclass | v2.1.3 |
| 3d-dna | v180922 | RepeatMasker | v4.1.2-p1 |
| Juicebox | v1.11.08 | HiSat2 | v2.2.1 |
| Minimap2 | v2.24-r1122 | Braker | v2.1.6 |
| Samtools | v1.10 | Gffread | v0.12.7 |
| Integrative Genome Viewer | v2.11.2 | SignalP | v6.0 |
| Tapestry | v1.0.0 | BUSCO | v5.3.2 |
| Blobtools | v1.1.1 | R | v4.2.0 |
| BLAST | 2.13.0+ | Circilize | v0.4.14 |
| Kmer Analysis Toolkit | V2.4.2 | InterProScan | v5.61 |
| Mummer | C3.23 |  |  |

## References

1. Abrusán, G., Grundmann, N., DeMester, L., and Makalowski, W. (2009). TEclass--a tool for automated classification of unknown eukaryotic transposable elements. Bioinformatics 25, 1329–1330.

2. Blanc, G., Agarkova, I., Grimwood, J., Kuo, A., Brueggeman, A., Dunigan, D.D., Gurnon, J., Ladunga, I., Lindquist, E., Lucas, S., et al. (2012). The genome of the polar eukaryotic microalga *Coccomyxa subellipsoidea* reveals traits of cold adaptation. Genome Biol 13, R39.

3. Camacho, C., Coulouris, G., Avagyan, V., Ma, N., Papadopoulos, J., Bealer, K., and Madden, T.L. (2009). BLAST+: architecture and applications. BMC Bioinformatics 10, 421.

4. Cao, S., Zhang, F., Zheng, H., Liu, C., Peng, F., and Zhou, Q. (2018a). *Coccomyxa antarctica* sp. nov. from the Antarctic lichen *Usnea aurantiacoatra*. PhytoKeys, 107-115.

5. Cao, S., Zhang, F., Zheng, H., Peng, F., Liu, C., and Zhou, Q. (2018b). *Coccomyxa greatwallensis* sp. nov. (Trebouxiophyceae, Chlorophyta), a lichen epiphytic alga from Fildes Peninsula, Antarctica. PhytoKeys, 39-50.

6. Danecek, P., Bonfield, J.K., Liddle, J., Marshall, J., Ohan, V., Pollard, M.O., Whitwham, A., Keane, T., McCarthy, S.A., Davies, R.M., et al. (2021). Twelve years of SAMtools and BCFtools. Gigascience 10.

7. Darienko, T., Gustavs, L., Eggert, A., Wolf, W., and Proschold, T. (2015). Evaluating the species boundaries of green microalgae (Coccomyxa, Trebouxiophyceae, Chlorophyta) using integrative taxonomy and DNA barcoding with further implications for the species identification in environmental samples. PLoS One 10, e0127838.

8. Davey, J.W., Davis, S.J., Mottram, C., and Ashton, P.D. (2020). Tapestry: validate and edit small eukaryotic genome assemblies with long reads. bioRxiv.

9. Dudchenko, O., Batra, S.S., Omer, A.D., Nyquist, S.K., Hoeger, M., Durand, N.C., Shamim, M.S., Machol, I., Lander, E.S., Aiden, A.P., et al. (2017). *De novo* assembly of the *Aedes aegypti* genome using Hi-C yields chromosome-length scaffolds. Science 356, 92–95.

10. Durand, N.C., Robinson, J.T., Shamim, M.S., Machol, I., Mesirov, J.P., Lander, E.S., and Aiden, E.L. (2016a). Juicebox provides a visualization system for Hi-C contact maps with unlimited zoom. Cell Syst 3, 99–101.

11. Durand, N.C., Shamim, M.S., Machol, I., Rao, S.S.P., Huntley, M.H., Lander, E.S., and Aiden, E.L. (2016b). Juicer provides a one-click system for analyzing loop-resolution Hi-C experiments. Cell Systems 3, 95–98.

12. Faluaburu, M.S., Nakai, R., Imura, S., and Naganuma, T. (2019). Phylotypic characterization of mycobionts and photobionts of rock tripe lichen in East Antarctica. Microorganisms 7.

13. Flynn, J.M., Hubley, R., Goubert, C., Rosen, J., Clark, A.G., Feschotte, C., and Smit, A.F. (2020). RepeatModeler2 for automated genomic discovery of transposable element families. Proc Natl Acad Sci U S A 117, 9451–9457.

14. Gray, A.P., Lucas, I.A.N., Seed, R., and Richardson, C.A. (1999). *Mytilus edulis chilensis* infested with *Coccomyxa parasitica* (Chlorococcales, Coccomyxaceae). Journal of Molluscan Studies 65, 289–294.

15. Greiner, S., Lehwark, P., and Bock, R. (2019). OrganellarGenomeDRAW (OGDRAW) version 1.3.1: expanded toolkit for the graphical visualization of organellar genomes. Nucleic Acids Res 47, W59–W64.

16. Gu, Z., Gu, L., Eils, R., Schlesner, M., and Brors, B. (2014). Circlize implements and enhances circular visualization in R. Bioinformatics 30, 2811–2812.

17. Gustavs, L., Schiefelbein, U., Darienko, T., and Pröschold, T. (2017). Symbioses of the green algal genera *Coccomyxa* and *Elliptochloris* (Trebouxiophyceae, Chlorophyta). In Algal and Cyanobacteria Symbioses, M. Grube, J. Seckbach, and L. Muggia, eds. (Europe: World Scientific), pp. 169–208.

18. Hoff, K.J., Lange, S., Lomsadze, A., Borodovsky, M., and Stanke, M. (2016). BRAKER1: Unsupervised RNA-Seq-based genome annotation with GeneMark-ET and AUGUSTUS. Bioinformatics 32, 767–769.

19. Jaag, O. (1933). *Coccomyxa* Schmidle-Monographie einer Algengattung (Bern: Gebrüder Fretz).

20. Kim, D., Paggi, J.M., Park, C., Bennett, C., and Salzberg, S.L. (2019). Graph-based genome alignment and genotyping with HISAT2 and HISAT-genotype. Nat Biotechnol 37, 907–915.

21. Koren, S., Walenz, B.P., Berlin, K., Miller, J.R., Bergman, N.H., and Phillippy, A.M. (2017). Canu: scalable and accurate long-read assembly via adaptive k-mer weighting and repeat separation. Genome Res 27, 722–736.

22. Kraege, A., Thomma, B.P.H.J., and Rovenich, H. (2023). Fastqc reports of sequencing data from *Coccomyxa viridis* SAG 216-4.

23. Kulichovà, J., Škaloud, P., and Neustupa, J. (2014). Molecular diveristy of green corticolous microalgae from two sub-Mediterranean European localities. European Journal of Phycology 49, 345–355.

24. Kurtz, S., Phillippy, A., Delcher, A.L., Smoot, M., Shumway, M., Antonescu, C., and Salzberg, S.L. (2004). Versatile and open software for comparing large genomes. Genome Biol 5, R12.

25. Laetsch, D.R., and Blaxter, M.L. (2017). BlobTools: Interrogation of genome assemblies. F1000 Research 6, 1287.

26. Lang, B.F., Beck, N., Prince, S., Sarrasin, M., Rioux, P., and Burger, G. (2023). Mitochondrial genome annotation with MFannot: a critical analysis of gene identification and gene model prediction. Front Plant Sci 14, 1222186.

27. Leliaert, F., Smith, D.R., Moreau, H., Herron, M.D., Verbruggen, H., Delwiche, C.F., and De Clerck, O. (2012). Phylogeny and molecular evolution of the green algae. Crit Rev Plant Sci 31, 1–46.

28. Li, H. (2021). New strategies to improve minimap2 alignment accuracy. Bioinformatics 37, 4572–4574.

29. Li, H., and Durbin, R. (2009). Fast and accurate short read alignment with Burrows-Wheeler transform. Bioinformatics 25, 1754–1760.

30. Li, L., Wang, S., Wang, H., Sahu, S.K., Marin, B., Li, H., Xu, Y., Liang, H., Li, Z., Cheng, S., et al. (2020). The genome of *Prasinoderma coloniale* unveils the existence of a third phylum within green plants. Nat Ecol Evol 4, 1220–1231.

31. Malavasi, V., Skaloud, P., Rindi, F., Tempesta, S., Paoletti, M., and Pasqualetti, M. (2016). DNA-Based taxonomy in ccologically versatile microalgae: A re-evaluation of the species concept within the coccoid green algal genus *Coccomyxa* (Trebouxiophyceae, Chlorophyta). PLoS One 11, e0151137.

32. Manni, M., Berkeley, M.R., Seppey, M., Simao, F.A., and Zdobnov, E.M. (2021). BUSCO update: novel and streamlined workflows along with broader and deeper phylogenetic coverage for scoring of eukaryotic, prokaryotic, and viral genomes. Mol Biol Evol 38, 4647–4654.

33. Mapleson, D., Accinelli, G.G., Kettleborough, G., Wright, J., and Clavijo, B.J. (2016). KAT: A K-mer Analysis Toolkit to quality control NGS datasets and genome assemblies. Bioinformatics 33, 574–576.

34. Marin, B. (2012). Nested in the Chlorellales or independent class? Phylogeny and classification of the Pedinophyceae (Viridiplantae) revealed by molecular phylogenetic analyses of complete nuclear and plastid-encoded rRNA operons. Protist 163, 778–805.

35. McFadden, G.I., and Melkonian, M. (1986). Use of Hepes buffer for microalgal culture media and fixation for electron microscopy. Phycologia 25, 551–557.

36. Morris, J.L., Puttick, M.N., Clark, J.W., Edwards, D., Kenrick, P., Pressel, S., Wellman, C.H., Yang, Z., Schneider, H., and Donoghue, P.C.J. (2018). The timescale of early land plant evolution. Proc Natl Acad Sci U S A 115, E2274–E2283.

37. Paysan-Lafosse, T., Blum, M., Chuguransky, S., Grego, T., Pinto, B.L., Salazar, G.A., Bileschi, M.L., Bork, P., Bridge, A., Colwell, L., et al. (2023). InterPro in 2022. Nucleic Acids Res 51, D418–D427.

38. Pertea, G., and Pertea, M. (2020). GFF utilities: GffRead and GffCompare. F1000 Research 9, 304.

39. Riehl, K., Riccio, C., Miska, E.A., and Hemberg, M. (2022). TransposonUltimate: software for transposon classification, annotation and detection. Nucleic Acids Res 50, e64.

40. Robinson, J.T., Thorvaldsdóttir, H., Winckler, W., Guttman, M., Lander, E.S., Getz, G., and Mesirov, J.P. (2011). Integrative genomics viewer. Nat Biotechnol 29, 24–26.

41. Schmidle, W. (1901). Über drei Algengenera. Berichte der deutschen botanischen Gesellschaft 19, 10–24.

42. Sciuto, K., Baldan, B., Maracto, S., and Moro, I. (2019). *Coccomyxa cimbria* sp. nov., a green microalga found in association with carnivorous plants of the genus *Drosera* L. European Journal of Phycology 54, 531–547.

43. Smit, A.F., Hubley, R., and Green, P. (2012). RepeatMasker (Retrieved from https://repeatmasker.org).

44. Sokolnikova, Y., Magarlamov, T., Stenkova, A., and Kumeiko, V. (2016). Permanent culture and parasitic impact of the microalga *Coccomyxa parasitica*, isolated from horse mussel *Modiolus kurilensis*. J Invertebr Pathol 140, 25–34.

45. Sokolnikova, Y., Tumas, A., Stenkova, A., Slatvinskaya, V., Magarlamov, T., and Smagina, E. (2022). Novel species of parasitic green microalgae *Coccomyxa veronica* sp. nov. infects *Anadara broughtonii* from Sea of Japan. Symbiosis 87, 293–305.

46. Štifterovà, A., and Neustupa, J. (2015). Community structure of corticolous microalgae within a single forest stand: evaluating the effects of bark surface pH and tree species. Fottea Olomouc 15, 113–122.

47. Tagirdzhanova, G., Scharnagl, K., Yan, X., and Talbot, N.J. (2023). Genomic analysis of *Coccomyxa viridis*, a common low-abundance alga associated with lichen symbioses. Sci Rep 13, 21285.

48. Teufel, F., Almagro Armenteros, J.J., Johansen, A.R., Gislason, M.H., Pihl, S.I., Tsirigos, K.D., Winther, O., Brunak, S., von Heijne, G., and Nielsen, H. (2022). SignalP 6.0 predicts all five types of signal peptides using protein language models. Nat Biotechnol 40, 1023–1025.

49. Tillich, M., Lehwark, P., Pellizzer, T., Ulbricht-Jones, E.S., Fischer, A., Bock, R., and Greiner, S. (2017). GeSeq - versatile and accurate annotation of organelle genomes. Nucleic Acids Res 45, W6–W11.

50. Trémouillaux-Guiller, J., Rohr, T., Rohr, R., and Huss, V.A.R. (2002). Discovery of an endophytic alga in *Ginkgo biloba*. Am J Bot 89, 727–733.

51. Vaschenko, M.A., Kovaleva, A.L., Syasina, I.G., and Kukhlevsky, A.D. (2013). Reproduction-related effects of green alga *Coccomyxa* sp. infestation in the horse mussel *Modiolus modiolus*. J Invertebr Pathol 113, 86–95.

52. Vaser, R., and Šikić, M. (2021). Time- and memory-efficient genome assembly with Raven. Nature Computational Science 1, 332–336.

53. Wick, R. (2018). Porechop (Retrieved from https://github.com/rrwick/Porechop).

54. Yahr, R., Florence, A., Škaloud, P., and Voytsekhovich, A. (2015). Molecular and morphological diversity in photobionts associated with *Micarea* s. str. (*Lecanorales*, Ascomycota). The Lichenologist 47, 403–414.

55. Zoller, S., and Lutzoni, F. (2003). Slow algae, fast fungi: exceptionally high nucleotide substitution rate differences between lichenized fungi Omphalina and their symbiotic green algae Coccomyxa. Mol Phylogenet Evol 29, 629–640.

